# Hippocampal Concentrations Drive Seizures in a Rat Model for Cefepime-induced Neurotoxicity

**DOI:** 10.1101/2022.05.26.493582

**Authors:** Emily Lesnicki, Gwendolyn M. Pais, Sylwia Marianski, Kimberly Valdez, Zoe Gibson, Jeffri Christopher, Marc H. Scheetz

## Abstract

**Background:** In high dose, cefepime causes neurotoxicity in patients with kidney injury; however, the relationship between exposure and observed neurotoxicity is not clear, and no animal model presently recapitulates the human condition.

**Objectives:** This study sought to describe plasma and tissue pharmacokinetics and pharmacodynamics (PK/PD) of cefepime in rats experiencing neurotoxicity.

**Methods:** Male Sprague-Dawley rats (n=21) received escalating cefepime total daily doses ranging from 531-1593 mg/kg body weight/day administered as a short infusion (0.5 mL/min) every 24h for 5 days. Cefepime was quantified in plasma, cerebral cortex and hippocampus via liquid chromatography-tandem mass spectrometry (LC-MS/MS). Multiple PK/PD models of cefepime transit between plasma and brain compartments (i.e. cerebral cortex and hippocampus) and neurotoxic response were explored using Monolix 2021R1 (LixoftPK).

**Results:** Exposure estimation of cerebral cortex demonstrated a median (IQR) AUC_0 –24_ and C_max 0 –24_ of 181.8 (85.2-661.3) mg · 24 h/liter and 13.9 (1.0-30.1) mg/L, respectively. The median cerebral cortex/blood percentage of penetration was 1.7%. Exposure estimation of hippocampus demonstrated a median (IQR) AUC_0 –24_ and C_max 0 –24_ of 291.4 (126.6-1091.6) mg · 24 h/liter and 8.8 (3.4-33.4) mg/L, respectively. The median hippocampus/blood percentage of penetration was 4.5%. Rats that reached a cefepime C_max_ of □17 mg/L in the hippocampus exhibited signs of neurotoxicity. A hippocampal cefepime concentration of 4.1 µg/100 mg brain tissue best described seizure stages >1 for cefepime-induced neurotoxicty.

**Conclusions:** A cefepime plasma AUC_0 –24_ of 28,000 mg•24h/L and hippocampal concentrations of 4.1 µg/100 mg brain tissue may be a threshold for cefepime-induced neurotoxicity. This model provides a methodology for future interrogation of the relationship between plasma concentrations, brain tissue concentrations, and neurotoxicity.

## Introduction

Antimicrobial resistance is a growing public health challenge, with a world-wide 4.95 million associated deaths associated in 2019, and 1.27 million deaths directly attributed to resistance^1^. For many antibiotics, standard recommended doses are no longer effective and as a result, clinicians have few options but to use maximal antibiotic exposure to treat resistant infections and cure infections. However, higher doses carry additional toxicological risks.

Cefepime is a broad-spectrum, cephalosporin used to treat bacterial infections such as pneumonia, urinary tract infections, and skin infections commonly caused by gram-positive and gram-negative bacteria^2^. It is the 4^th^ most commonly used Gram-negative antibiotic administered to hospital patients^3^. While a class effect of neurotoxicity is known for β-lactam agents such as cefepime^4^, cefepime in particular is associated with a high rate of neurotoxicity. A retrospective cohort study found a cumulative incidence of neurotoxicity of 41% with cefepime, 24% with meropenem, and 35% with piperacillin/tazobactam^5^. When comparing the convulsive activity of other β-lactams, cefepime is ∼1.6 fold more pro-convulsive than penicillin and ∼2x more than imipenem. Conversely, ceftriaxone is 10x less pro-convulsive than cefepime^6^. In 2012, The Food and Drug Administration issued a warning of seizure risk associated with cefepime use in patients suffering from renal impairment that do not receive appropriate dose adjustments^7^. Of the 59 individuals displaying neurotoxic outcomes, 58 of those patients had renal dysfunction, and 56 patients received a higher than recommended dose for their organ function. The safety of cefepime in certain patient populations has been routinely examined, especially after adverse effects have been observed at high rates within standard recommended dosing regimens.

### Clinical Pharmacology

The standard adult dose of cefepime is 2 grams every 8 hours via intravenous infusion over 30 minutes for common Gram-negative pathogens. The drug follows “linear” elimination kinetics; it has an observed half-life of 2 (±0.3) hours and a total body clearance of 120 (±8) mL/min in healthy adults^2^. Cefepime is primarily excreted by the kidneys, therefore patients with reduced renal function are more susceptible to increased exposures if doses are not decreased. Notably, kidney damage is the most common comorbidity among those suffering from cefepime neurotoxicity. Patients with kidney disease can have the cefepime half-life increase to 13 hours compared to patients with normal clearance^8^. The average age of patients suffering from cefepime neurotoxicity was 67 years old, equally affecting men (49%) and women (51%)^9^. The most observed clinical manifestations of cefepime neurotoxicity include loss of consciousness, aphasia, confusion, non-convulsive status epilepticus (NCSE), encephalopathy, seizure disorders, myoclonus, and other neuropsychiatric symptoms^10, 11^. In patients with NCSE, electroencephalographic (EEG) methods were used to observe altered brain activity, which showed tri-phasic, generalized slow, and multi-focal sharp waves, all of which are abnormal^12^.

Roughly 86% of the cefepime is recovered in the urine unchanged in patients with normal renal function^13^. In addition to renal excretion, cefepime is metabolized into N-methylpyrrolidine (NMP) and further into NMP-N-oxide and an epimer of cefepime. Studies suggest that penicillin related compounds are actively transported across the blood-brain-barrier^14^. Cefepime can penetrate the cerebral spinal fluid (CSF) with observed median CSF-plasma concentration ratios of 19%, as demonstrated in a pharmacokinetic rat model^15^. These results are in agreement with other animal studies^16,17^, as well as transit in humans^18^.

When the blood-brain-barrier permeability is disrupted, greater concentrations of the drug are likely to reach the brain, but especially as the degree of renal failure increases. Increased central nervous system (CNS) penetration has also been observed in patients with sepsis, CNS infection, and brain injury^19^. While there are data on penetration of cefepime in the CSF and plasma, little is known about accumulation in the brain and relationship with toxicity. The accumulation of cefepime in the brain may be the important driving factor linking cefepime and neurotoxicity.

The specific therapeutic plasma concentrations that define cefepime neurotoxicity are not clear; though, some have suggested trough concentrations > 22 mg/L^20^; however, the precision of trough concentrations to predict neurotoxicity may not be accurate. The estimated mean probability of neurotoxicity at T_>22_ in one study was 51.4%, which is a rate far beyond what is seen in clinical practice^21^. Such a model could be used to simulate the human toxicity threshold as there is no threshold goal to date. The objectives of the current research are to gain a better understanding of the pharmacokinetic exposures resulting in neurotoxic endpoints.

### Mechanism of Neurotoxicity

Although the mechanism contributing to cefepime induced neurotoxicity is not entirely understood, studies show that the adverse events may be at least partially mediated by cefepime binding to the gamma-aminobutyric acid (GABA_A_) receptor^22, 23^. Cefepime demonstrates a high binding affinity and binds competitively to the GABA_A_ subtype receptors in a concentration dependent manner^22^. The inhibition of GABA receptor activation causes hyperexcitability, resulting in a lower seizure threshold^22^. However, other potential mechanisms of cefepime neurotoxicity likely exist to explain the higher rates of neurotoxicity.

### Models of Neurotoxicity

Rodent models are routinely used to assess potential convulsive risk of β-lactam antibiotics. Researchers have administered various cephalosporins to test the range of convulsive effects of β-lactam antibiotics. Several models of cefepime neurotoxicity have been established such as the PTZ method to chemically induce seizures by acting as a GABA inhibitor and electroconvulsive shock in corneal kindled mice to determine the convulsive liability of cefepime. The neurotoxicity outcome is quickly achieved by lowering the seizure threshold^24^. Similarly, intracerebral administration of cefepime also produces robust seizure responses within minutes^23^. A gap in literature is a clinically relevant animal model to define the systemic pharmacokinetic exposure that results in neurotoxicity, specifically in the context of renal impairment. The rodent model has high translational capacity due to similar brain structure and neurotransmitters, which is why it is used in various seizure models. However, a rodent model for neurotoxicity that delivers cefepime systemically does not yet exist.

Apart from renal impairment, brain injury or neurological disorders may also be risk factors for convulsive activity. Epilepsy can lower the seizure threshold, increasing the risk of cefepime-induced convulsions^24^. The symptoms consistent with neurotoxicity in these animal models include rolling, wild running, clonic convulsions, falling down, clonus of the forelimbs, and death^22,23^.

## Materials and Methods

### Experimental design and animals

The animal toxicology study was conducted at Midwestern University (IACUC 2793). Male Sprague-Dawley rats (mean weight 260-300 g) were obtained from Envigo (Indianapolis, IN, USA).

### Chemicals and reagents

Animals were administered clinical grade cefepime hydrochloride for injection (Apotex Corporation, Weston, FL, USA). Normal saline (Abbott Laboratories, Chicago, IL, USA) and heparin (Covetrus, Portland, ME, USA) were used in sampling methods. Folic acid (Sigma-Aldrich, St. Louis, MO, USA) dissolved in 0.3 mmol/L NaHCO_3_ (VWR, Radnor, PA, USA) was used.

Analytical grade cefepime hydrochloride (Apotex Corporation, Weston, FL, USA) and ceftazidime (Acros Organics, NJ, USA) pentahydrate as an internal standard were used for liquid chromatography-tandem mass spectrometry (LC-MS/MS) assays. Milli-Q water was obtained from Aqua Solutions purified water dispensing system at Midwestern University. LC–MS/MS grade acetonitrile (VWR, Radnor, PA, USA), formic acid (VWR, Radnor, PA, USA), methanol (VWR, Radnor, PA, USA) and frozen, non-medicated, non-immunized, pooled Sprague–Dawley rat EDTA plasma (BioIVT, Westbury, NY, USA) were used to generate standard curves.

### Drug Administration

Animals were temporarily anesthetized with 5% isoflurane via calibrated vaporizer with a charcoal canister and maintained on 2-3% isoflurane via nasal cone for approximately 3 minutes until all folic acid was administered intraperitoneally (IP) (divided into one or two doses). Cefepime was administered via single jugular vein catheter 30 minutes post folic acid.

### Experimental Protocol

#### Maximum human dosing pre-and post-acute kidney injury (AKI) (Aim 1)

Rats received 531 mg/kg/day of cefepime (the allometric scaled dose from maximum human dosing is 86 mg /kg/day based on a package insert dose of 2000 mg three times daily for a 70 kg patient)^2^ over approximately 2 minutes via jugular vein catheter. Folic acid (250 mg/kg) was administered IP in the hind limb while under isoflurane. Animals served as their own controls. Cefepime was dosed once daily and plasma samples were collected at various times before and after folic acid administration to mimic pre and post-AKI conditions.

#### Maximum tolerated dosing (Aim 2)

Rats received increasing doses of cefepime ranging from 500-2000 mg/kg/day as IV infusions given over approximately 2 minutes to determine the maximum tolerated dose (MTD)^24^. Folic acid (250 mg/kg) was administered IP on the first day prior to cefepime. The second group received the MTD of cefepime as a 0.5 mL/min infusion. Animals were observed for neurotoxic outcomes. Convulsive behavior was visually assessed according to the following modified Racine scale^25^: stage 0 no response, stage 1 ear and facial twitching, stage 2 myoclonic body jerks, stage 3 forelimb clonus and rearing, stage 4 clonic convulsions and turned on the side, stage 5 generalized clonic convulsions and turned on the back, stage 6 status epilepticus, and stage 7 loss of life. EEG activity was recorded to observe cefepime signature of non-convulsive status epilepticus. Sampling procedures were repeated as previously described.

#### Seizure characterization after MTD cefepime exposures (Aim 3)

Rats received 250 mg/kg folic acid IP in two divided doses, dissolved in 0.3 mmol/L sodium bicarbonate on the first day of protocol and received 100 mg/kg folic acid each there thereafter to reduce renal function and slow cefepime clearance. Rats received the MTD (either 1593 or 1250 mg/kg) of cefepime. Animals were placed in metabolic cages each day, and urine was collected over 24-hour periods over a period of 5 days. The experimental design is outlined in Figure 1. Convulsive behavior was assessed by a modified Racine scale^25^ as previously described. On the final day, rats were anesthetized with 100mg/kg ketamine and 10 mg/kg xylazine IP. Terminal plasma and serum (1 mL) were collected. Tissues were perfused with chilled saline to prevent contamination with circulating blood and brains were harvested. Serial sacrifice occurred before or during convulsive episodes as outlined in Table 1. Seizure stages were determined by the last observed seizure before brain harvest. Concentrations of cefepime are expressed as µg/100 mg of brain tissue.

**Table 1.**
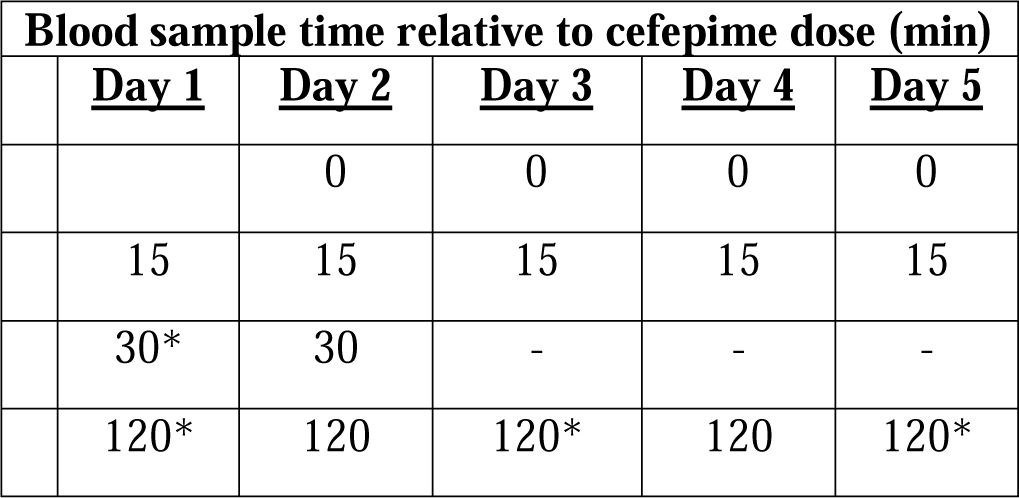
Sampling and serial sacrifice schedule. Sacrifice denoted by *.

**Figure 1.**
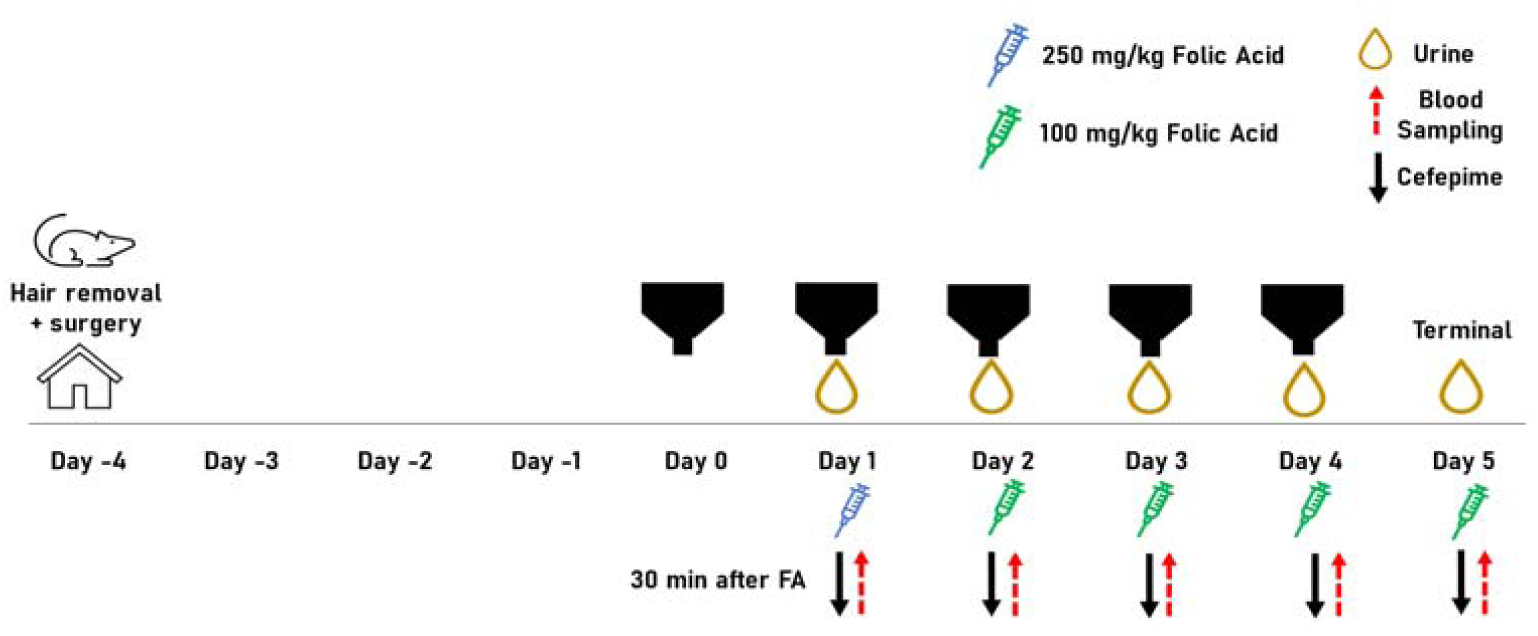
Experimental design timeline (Aim 3)

### Blood collection

Double jugular vein cannulation surgery was performed, and animals were allowed to recover four days prior to first day of sampling. Blood samples were drawn from a single jugular vein catheter in a sedation free manner when possible and collected into EDTA tubes. Blood samples of 0.15 mL were replaced with equivalent volume of normal saline to maintain euvolemia. Dilute heparin (0.1 mL) was administered to prevent clotting. Samples were taken at various time points (i.e. 15, 30, 60, and 120 minutes). Blood was centrifuged for 10 minutes at 3,000 g, plasma supernatant was collected, then stored at −80°C until analysis.

### Plasma Analysis

Due to high cefepime concentrations in the plasma, samples were diluted (i.e.136x, 62x, 32x, 8x, or 4x) with corresponding matrix so concentrations were within standard curve range. Standard curves were prepared using fresh cefepime and ceftazidime. Plasma samples volumes of 40 µl were combined with 4 µl of internal standard (10 µg/ml ceftazidime) and subject to protein precipitation using 456 µl methanol and 1% formic acid. Samples were centrifuged at 16,000g for 10 minutes at 4°C and 100 µl supernatant was collected for analysis. The plasma concentrations were quantified by LC-MS/MS using standard curves for each matrix. Milli-Q water containing 0.1% formic acid and acetonitrile (flow rate of 0.5 ml/min) are used as aqueous (A) and organic (B) solvents, respectively, at the following ramping transitions: 0.00 min A (90%) →B (10%), 1.50 min A (90%) →B (10%), 2.50 min A (10%) →B (90%), 5.40 min, A (10%) →B (90%), 5.50 min A (90%) →B (10%), and 10 min A (90%)→B (10%). A Waters (2.1×100mm, 1.7μm) Acquity UPLC CSH C18 column (Agilent Technologies, Inc., Santa Clara, CA, USA) was utilized.

### Tissue Homogenization

A BeadBug (Benchmark Scientific, Sayreville, NJ, USA) tissue homogenizer was used to homogenize cerebral cortex and hippocampus samples. Brain samples were first manually cut up into smaller pieces using dissecting scissors. Approximately 100 mg of cortex and hippocampus were placed into screw caps with a three-fold volume of sample weight of MilliQ, and Zirconium mm beads. Tissues underwent 3 cycles at 2500 rpm for 30 seconds with a 30 second rest interval repeated twice. Assays were prepared with brain homogenates using 40 µl sample, 4µl of ceftazidime internal standard, and 456 µl of 0.1% formic acid in methanol. Samples were centrifuged at 16000 g for 10 minutes at 4°C.

### Pharmacokinetic Modeling

Plasma concentrations from Aim 1 experiments were used to run a one-compartmental linear elimination infusion model in Monolix. This model has fit the data well in preliminary studies. Clearance and volume of distribution parameters of cefepime were obtained from the fitted model and used to calculate half-life before and after kidney injury.

Plasma and tissue concentrations used to develop a physiologically based PK model in Monolix 2021R1 (LixoftPK). Animals that were subjected to similar experimental protocols were used in the final model. Multiple PK/PD models of cefepime transit between plasma and brain compartments (i.e. cerebral cortex and hippocampus) and neurotoxic response were explored using Monolix. PK parameters and exposures during the first 24 hours (i.e., area under the concentration-time curve from 0 to 24 h [AUC_0 –24_] and maximum concentration of drug in plasma from 0 to 24 h [C_max0 –24_]) were calculated from Empiric Bayes Estimated concentrations given exact dosing schedules for each rat in Simulx (Lixoft). PK parameters from brain tissue were correlated with convulsive behavioral scores as described by a modified Racine scale^25^.

### Statistical Analysis

All statistical analyses and graphics were generated using GraphPad Prism 9. Mean half-life of cefepime in pre and post-AKI conditions was analyzed by paired t-test. Analyses comparing mean cefepime concentrations and PK parameters in cortex and hippocampus of seizures stage groups (≤ 1 or >1), and differences in each cohort were done by independent t-test. All tests were two-tailed with statistical significance set at alpha 0.05.

## RESULTS

### Folic acid-induced AKI

Three animals were included in the initial pharmacokinetic analysis. A one-compartment linear elimination model fit the data well for plasma (R^2^=0.92). Mean half-life of cefepime for the pre-AKI condition was 0.382 hours and post-AKI mean half-life was 4.27 hours. The mean elimination half-life of cefepime increased with the presence of AKI, but differences pre and post folic acid treatment were not statistically significant; however, power was constrained (Figure 2). As these data were utilized to determine if kidney function was impacted with a new method of experimental kidney injury (i.e. folic acid), it was deemed that folic acid resulted in impairment of kidney function by a factor of ∼10.

**Figure 2.**
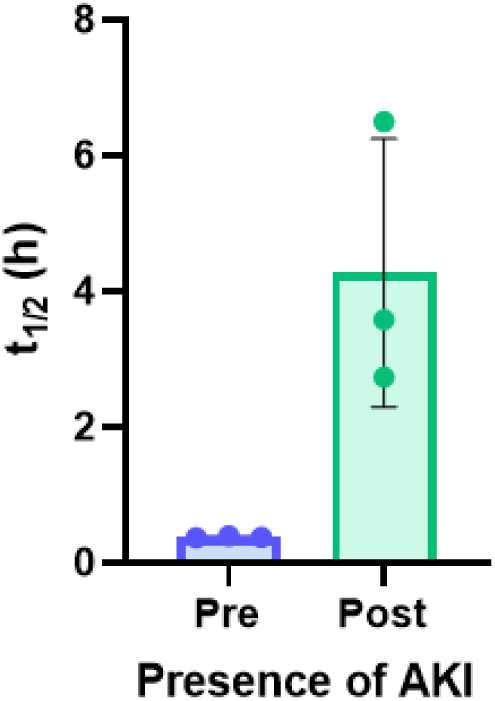
Impact of folic acid-induced AKI on the elimination half-life of cefepime. Values are expressed as mean ± SD, n=3 rats before and after folic acid administration. Pre AKI mean half-life is 0.382 hours and post mean half-life is 4.27 hours. Cefepime half-life was not significantly altered after folic acid (p=0.08 by paired t-test).

### Cefepime accumulation in the brain

Cefepime concentrations in the cerebral cortex and hippocampus were significantly higher in rats exhibiting seizure stages >1 (5.775 ± 2.71 µg/100 mg cortex tissue vs 1.55 ± 0.94 µg/100 mg cortex tissue, p <0.0003, and 6.85 ± 2.583 µg/100 mg hippocampal tissue vs 1.38 ± 0.76 µg/ 100 mg hippocampal tissue, p<0.0001). Cefepime concentrations in the hippocampus demonstrated a clear cut-off at 4.1 µg/100 mg hippocampal tissue, for cefepime-induced neurotoxicity (Figure 4).

**Figure 3:**
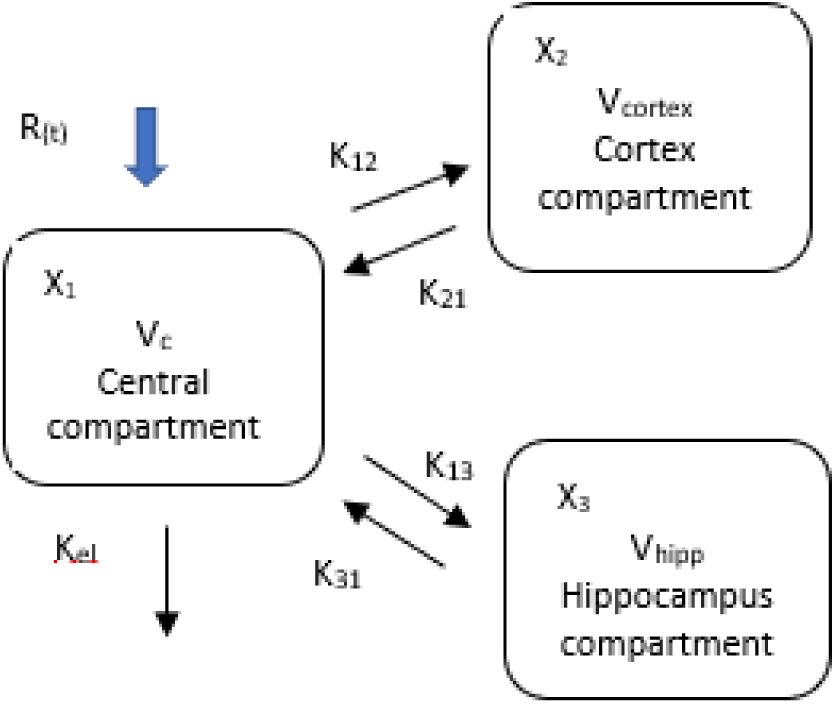
Schematic of three-compartmental PK model. Abbreviations: PK, pharmacokinetic; R(t), dose administration rate; K_el_, elimination rate constant; V_c_, volume of central compartment; V_cortex_, volume of cerebral cortex compartment; V_hipp_, volume of hippocampus compartment; K_12_, rate constant to cortex from central compartment; K_21_, rate constant to central from cortex compartment; K_13_, rate constant to hippocampus compartment from central compartment; K_31_, rate constant to central compartment from hippocampus compartment; X_1_, amount in central compartment; X_2_, amount in cortex compartment; X_3_, amount in hippocampus compartment.

**Figure 4.**
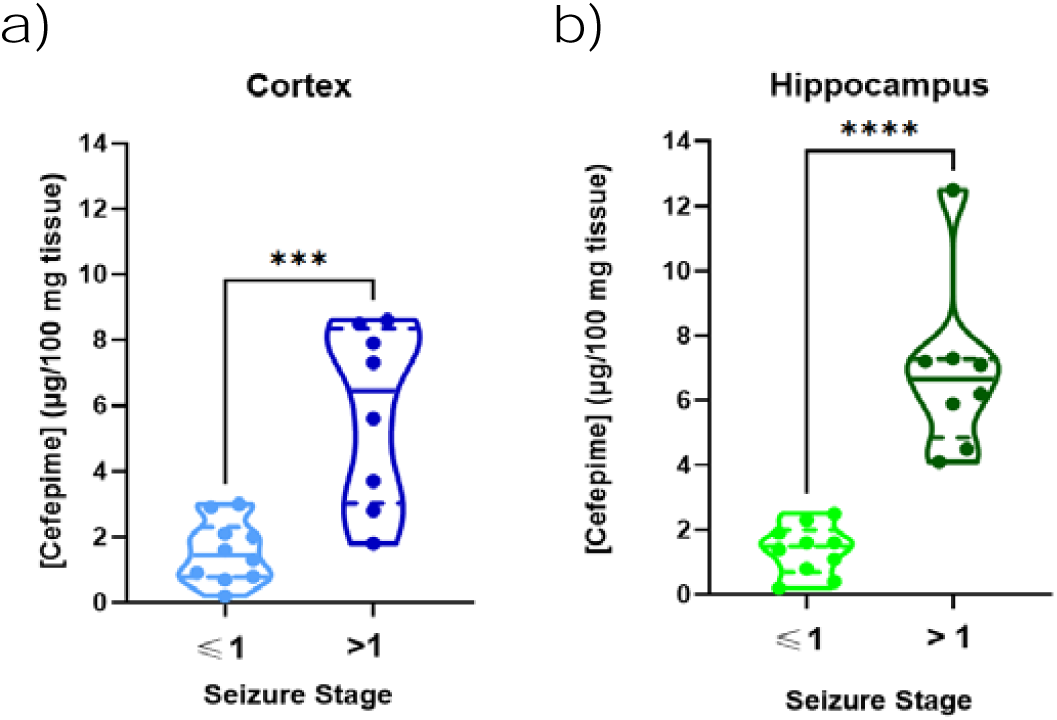
Concentrations of cefepime in the rat hippocampus and cerebral cortex relative to seizure stage. Cefepime concentrations expressed as µg/100 mg brain tissue. Concentrations in the (a) cerebral cortex and (b) hippocampus were higher in rats exhibiting seizure stages >1 (p = 0.0003 and p<0.0001, respectively, by student’s t-test) compared to rats exhibiting seizure stages ≤ 1.

### Cefepime pharmacokinetic model

A total of 21 rats received cefepime and contributed PK data. All available plasma samples that were collected were used in model building and analysis. The final model was a three-compartmental model for plasma PK, cerebral cortex, and hippocampus (Figure 3). The median parameter values (with the coefficient of variation percentage [CV%]) for the rate constants to the cerebral cortex from the central compartment (K_12_), to the central compartment from the cerebral cortex compartment (K_21_), to the hippocampus from the central compartment (K_13_), to the central compartment from the hippocampus compartment (K_31_), were 1.01 h^-1^, 1.98 h^-1^ [6.38%], 0.15 h^-1^ [26.9%], and 0.2 h^-1^ [46.5%], respectively. The model fit the data well for plasma with predictive performance of coefficients of determination (R^2^) were Bayesian [R^2^ = 0.60] for plasma, Bayesian [R^2^ = 0.99] for cerebral cortex, and Bayesian [R^2^ = 0.98] for hippocampus (Figure 6).

**Figure 5.**
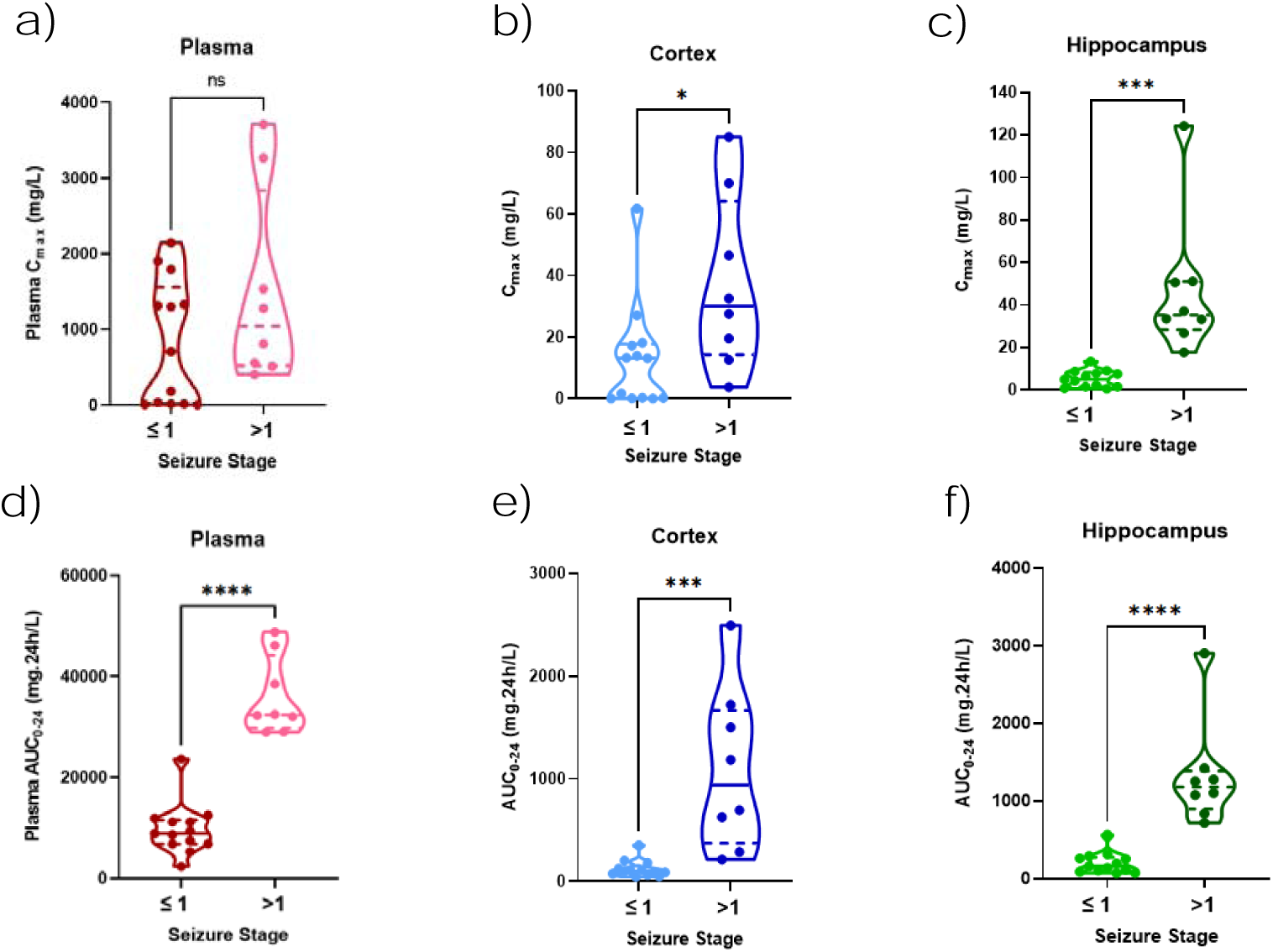
PK parameters and exposures during the first 24h calculated from Empiric Bayes Estimated concentrations estimated every 0.1 hours given exact dosing schedules for each rat. Cefepime Cmax_0 –24_ concentrations in the (b) cerebral cortex and (c) hippocampus were significantly higher in rats exhibiting seizure stages >1 (p = 0.023 and p = 0.0002, respectively). Cefepime AUC_0 –24_ (mg·24/L) in (d) plasma (p <0.0001), (e) cerebral cortex (p = 0.0003), and (f) hippocampus (p <0.0001) were significantly higher in rats exhibiting seizure stages >1.

**Figure 6.**
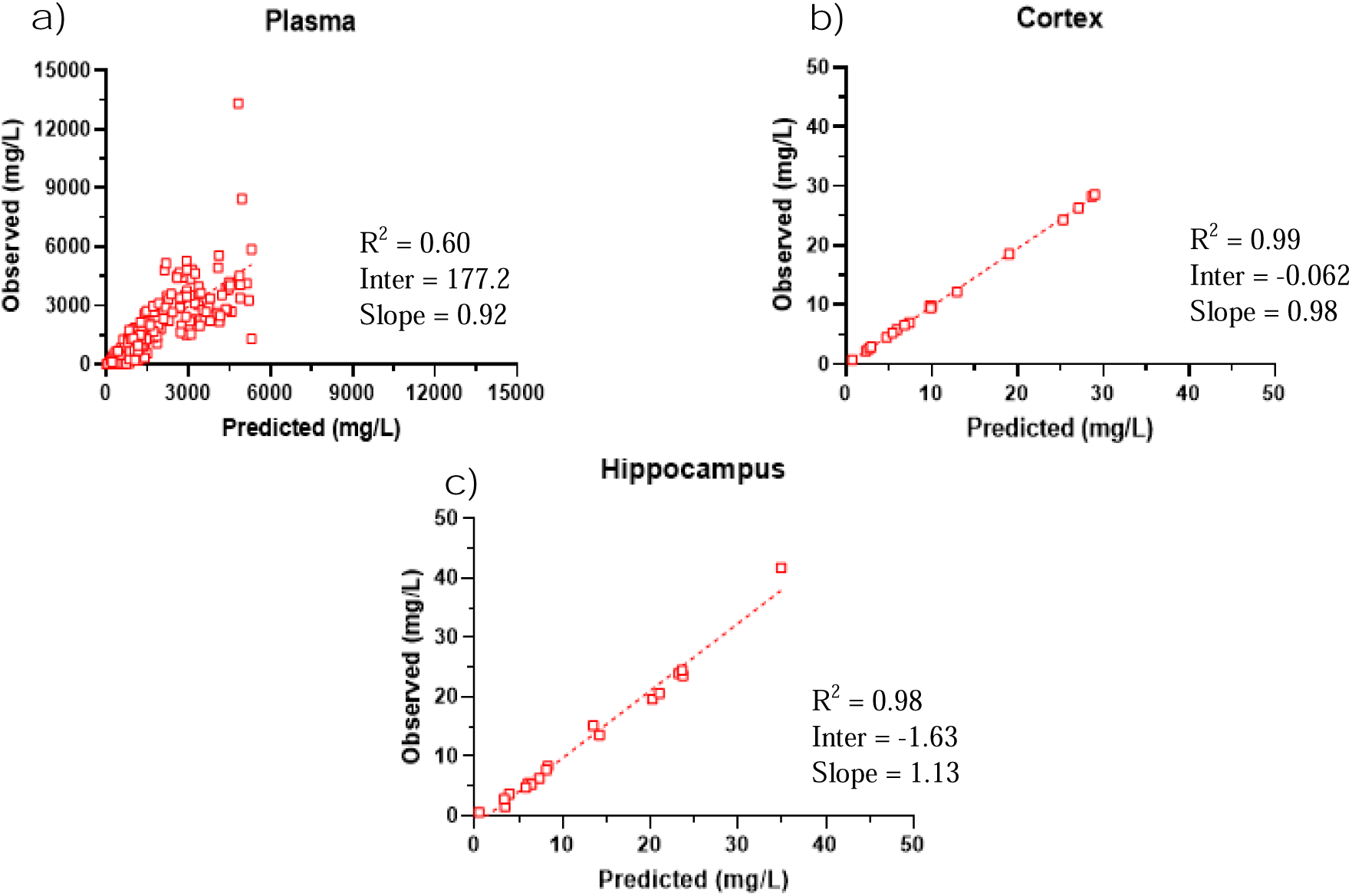
Observed versus predicted Bayesian plots from the [final] model for (a) plasma, (b) cerebral cortex, and (c) hippocampus.

### Cefepime pharmacokinetic exposures and percent penetration

Exposure estimation revealed a plasma median [IQR] half-life, AUC_0–24_, and C_max 0–24_, of 2.2 (1.1-5.8) h, 11916.5 (8060.5-32192.5) mg · 24 h/liter, and 809.4 (110.7-1664.8) mg/L from the first dose, respectively. Exposure estimation of cerebral cortex demonstrated a median [IQR] AUC_0–24_ and C_max 0 –24_ of 181.8 (85.2-661.8) mg · 24 h/liter and 13.9 (1.0-30.1) mg/L, respectively. The median cerebral cortex/blood percentage of penetration was 1.7%. Exposure estimation of hippocampus demonstrated a median [IQR] AUC_0–24_ and C_max 0 –24_ of 291.4 (126.6-1091.6) mg · 24 h/liter and 8.8 (3.4-33.4) mg/L, respectively. The median hippocampus/blood percentage of penetration was 4.5%. PK exposures for the first 24 h described in Figure 5. The complete pharmacokinetic exposures and percentages of cefepime penetration for all animals are summarized in Tables 3 and 4.

**Table 2:** Parameter values from preliminary AKI model

| RAT ID | OCCASION | V | Cl | $K_{el}$ | $t_{1/2}$ Pre | $t_{1/2}$ Post |
| --- | --- | --- | --- | --- | --- | --- |
| 35 | 1 | 0.15 | 0.28 | 1.87 | 0.37 |  |
| 35 | 2 | 0.15 | 0.038 | 0.25 |  | 2.74 |
| 36 | 1 | 0.093 | 0.17 | 1.83 | 0.38 |  |
| 36 | 2 | 0.093 | 0.018 | 0.19 |  | 3.58 |
| 37 | 1 | 0.092 | 0.16 | 1.74 | 0.398 |  |
| 37 | 2 | 0.092 | 0.0098 | 0.11 |  | 6.51 |

Pharmacokinetic parameters estimated by Monolix (Lixoft) software for a one-compartment linear elimination model. Occasion defined as pre and post AKI. Abbreviations: AKI, acute kidney injury; V, volume of distribution; Cl, clearance of elimination; $K_{el}$ , elimination constants (where $K_{el} = Cl/V$ ); $t_{1/2}$ , elimination half-life (where $t_{1/2} = 0.693/K_{el}$ ).

**Table 3.** Cefepime plasma, cerebral cortex, and hippocampus PK exposures estimated using Bayesian posteriors for AUC_0 –24_ and C_max0 –24_

| Animal | $\text{AUC}_{0-24}$<br>($\text{mg}\cdot\text{h}/\text{L}$)<br>plasma | $\text{C}_{\text{max}0-24}$<br>($\text{mg}/\text{L}$)<br>plasma | $\text{AUC}_{0-24}$<br>($\text{mg}\cdot\text{h}/\text{L}$)<br>cortex | $\text{C}_{\text{max}0-24}$<br>($\text{mg}/\text{L}$)<br>cortex | $\text{AUC}_{0-24}$<br>($\text{mg}\cdot\text{h}/\text{L}$)<br>hippocampus | $\text{C}_{\text{max}0-24}$<br>($\text{mg}/\text{L}$)<br>hippocampus | $t_{1/2}$<br>(h) |
| --- | --- | --- | --- | --- | --- | --- | --- |
| 35 | 2349.81 | 707.69 | 45.99 | 13.16 | 74.081 | 4.38 | 0.9 |
| 36 | 5293.99 | 1329.18 | 64.97 | 13.94 | 105.050 | 4.98 | 1.31 |
| 37 | 8646.59 | 1301.66 | 125.75 | 17.23 | 196.13 | 9.13 | 3.15 |
| 45 | 48805 | 1281.26 | 1723.33 | 46.54 | 1425.84 | 51.17 | 13.86 |
| 46 | 38542.1 | 809.39 | 1502.36 | 32.59 | 1106.01 | 33.55 | 9.36 |
| 47 | 32329.3 | 512.88 | 1186.02 | 19.55 | 1077.18 | 26.82 | 6.3 |
| 48 | 11138.1 | 1313.82 | 75.40 | 13.26 | 71.029 | 6.44 | 0.77 |
| 49 | 32055.6 | 559.53 | 695.51 | 12.59 | 1253.48 | 33.32 | 5.33 |
| 50 | 8934.17 | 14.99 | 181.83 | 0.33 | 254.15 | 1.59 | 1.155 |
| 51 | 29037.3 | 3712.45 | 628.18 | 69.97 | 1278.2 | 50.67 | 4.076 |
| 52 | 6846 | 1903.06 | 90.06 | 27.050 | 114.67 | 7.53 | 0.72 |
| 53 | 11164.8 | 2146.97 | 353.93 | 61.74 | 316.04 | 13.43 | 1.78 |
| 54 | 11916.5 | 1793.33 | 80.48 | 18.19 | 90.205 | 7.86 | 0.7 |
| 55 | 28971.7 | 3269.52 | 214.79 | 27.55 | 718.72 | 37.15 | 2.77 |
| 56 | 46125.5 | 1536.19 | 2493.59 | 84.99 | 2905.98 | 124.19 | 40.76 |
| 57 | 9380.96 | 15.08 | 50.35 | 0.087 | 261.72 | 1.59 | 1.78 |
| 58 | 23682.4 | 187.26 | 204.04 | 1.70 | 565.96 | 8.80 | 4.076 |
| 59 | 12505.4 | 34.24 | 101.71 | 0.30 | 291.36 | 2.37 | 2.24 |
| 60 | 32517.7 | 406.12 | 288.93 | 3.77 | 838.37 | 17.70 | 6.3 |
| 61 | 7474.34 | 8.19 | 111.745 | 0.13 | 160.76 | 0.80 | 1.36 |
| 62 | 6884.11 | 6.43 | 113.4 | 0.12 | 138.50 | 0.64 | 0.98 |
| **Median** | 11916.5 | 809.4 | 181.8 | 13.9 | 291.4 | 8.8 | 2.2 |
| **(IQR)** | 8060.5-32192.5 | 110.7-1664.8 | 85.3-661.8 | 1.0-30.1 | 126.6-1091.6 | 3.4-33.4 | 1.1-5.8 |

Abbreviations: $C_{\max 0-24}$ , maximum concentration at 24 h; $AUC_{0-24}$ , area under the curve at 24 h; $t_{1/2}$ , half-life; IQR, interquartile range

**Table 4.**
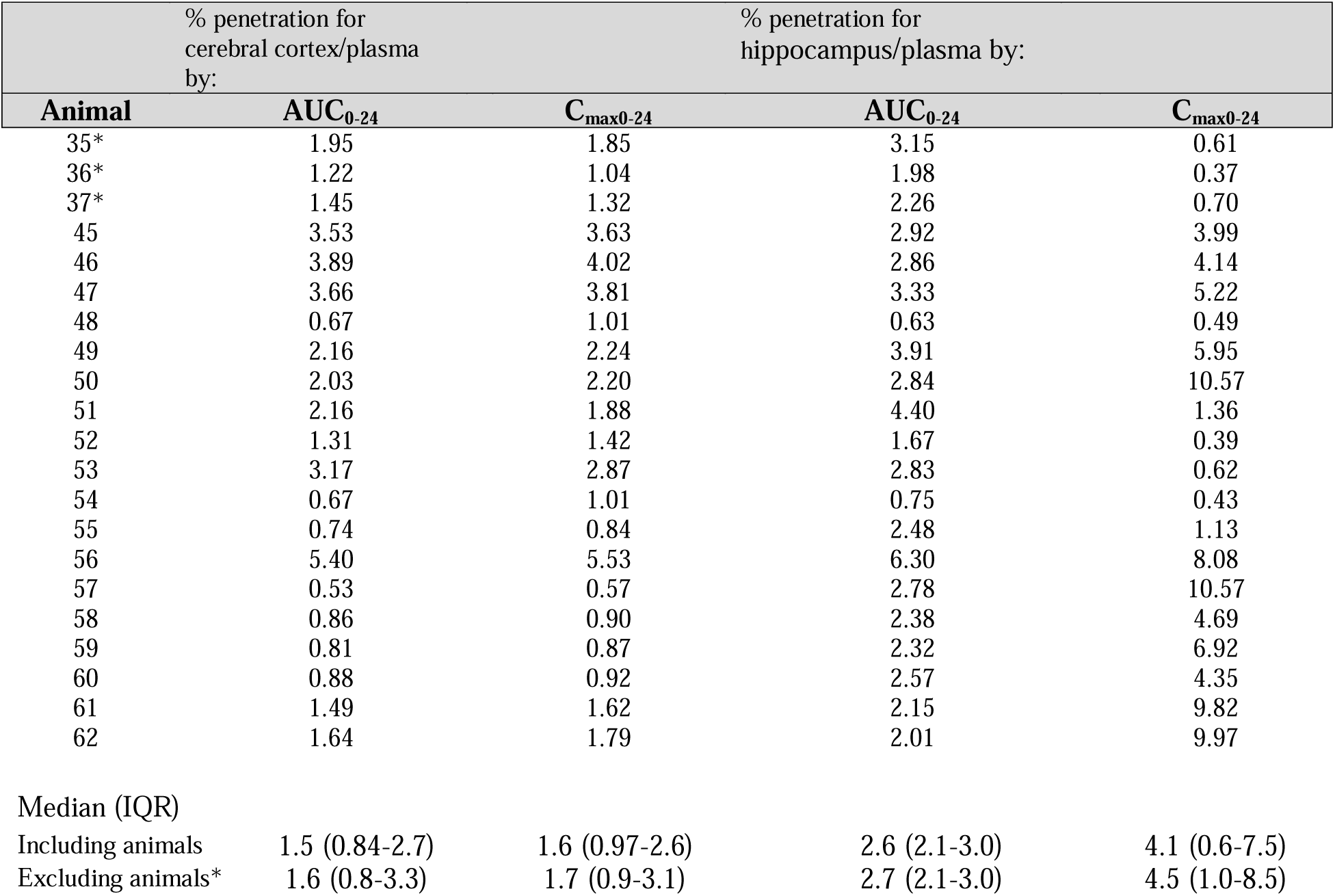
Percent of cefepime penetration Abbreviations: C_max 0 –24_, maximum concentration at 24 h; AUC _0 –24_, area under the curve at 24 h; t_1/2_, half-life; IQR, interquartile range. *No cerebral cortex or hippocampus samples were collected in these animals.

## Discussion

We induced kidney injury in rats and identified important exposure response relationships between cefepime and seizure stage. Hippocampal concentrations best described seizure stages >1 for cefepime-induced neurotoxicity. Previous research has demonstrated that when cefepime is administered intracerebrally, seizure responses are robust^23^, however, was unclear how much of the drug gets into the brain when administered systemically. We found that a cefepime plasma AUC_0 –24_ around 28,000 mg·24h/L corresponded to a hippocampal concentration of 4.1 µg/100 mg brain tissue in animals exhibiting greater seizure activity (seizure stages >1). The estimated C_max0 –24_ exposure was significantly higher for animals experiencing neurotoxic outcomes in the cortex and the hippocampus. These effects were most apparent in the hippocampal analysis, suggesting that hippocampal cefepime concentrations are responsible for driving seizures. The corresponding AUC_0-24_ cefepime plasma and cortex may also be linked to seizure outcome. Future work will be required to better understand the full relationships between plasma concentrations, various brain tissue concentrations, and neurotoxicity.

In our study we found a greater median hippocampus/blood percent penetration of 4.5% by C_max0 –24_, which was greater than the cerebral cortex/ blood penetration. Previous PK rat models showed that the median CSF/blood percentage of penetration of cefepime by AUC_0 –24_ was 19% and 3% by C_max0 –24_ ^15^. Similarly, other animal studies evaluated the transit of cefepime to the target areas have demonstrated cefepime CSF concentrations between 16.2 and 36%^16,17^. The data are also consistent with findings in human subjects, which found a percent penetration of 23% to the CSF^18^. Brain concentration findings in humans are more rare as tissue is difficult to obtain. However, microdialysis studies with other β-lactams demonstrate that brain concentrations are in line with class effects^26^. The PK of the parent compound has been evaluated extensively, but future studies should also consider the PK of the metabolites.

Our research identified that rats did not become neurotoxic in the absence of kidney injury in preliminary experiments. Renal impairment is a known risk factor of cefepime neurotoxicity, and the half-life of cefepime increased when AKI was present. Large doses of folic acid causes crystallization of the proximal tubule in the rat and has demonstrated to be an effective method to recreate the condition in the animal model.

PK-PD modeling has been useful in early stages of drug development and is an important tool for determining the efficacy and safety of a drug. By doing so, we can better understand toxicity outcomes and define the thresholds for toxicity. Animal models are frequently used in the PK evaluation of antimicrobial therapies. Although the PK of cefepime has been defined, the full PK/PD drivers are not well understood. This is the first study that quantitatively describes the transit of cefepime from the plasma to the cerebral cortex and hippocampal brain regions in rats experiencing neurotoxicity. This systemic exposure model is clinically relevant as the rat PK/PD model can be used to simulate the human toxicity threshold.

The mechanism for the CNS effects of cefepime remains unclear. The proposed explanation is attributed to its ability to cross the blood–brain barrier to bind competitively to the GABAergic receptor to suppress inhibitory neurotransmission^22^. Another suggested pathophysiology of these effects is a dysregulated lipid metabolism. Because the brain is a lipid-rich organ, dysregulated homeostasis may contribute to the development of cefepime neurotoxicity^27^. Cefepime has been found to dysregulate the glycerophospholipid profile in the corpus striatum in mice receiving intraperitoneal injection. The number of dysregulated lipids increased after 5 days of exposure and changes in composition and structure were also observed. Moreover, the proportion of GABAergic neurons are high in the cortex and hippocampus but may be higher within the striatum^27^. This area may be more sensitive to cefepime treatment. Our study did not have adequate brain samples to isolate and analyze the corpus striatum. Further studies are warranted.

There are several limitations in this study. Some animals did not contribute complete data; however, all available samples were used to inform the model. Also, experimental protocols differed slightly for the various studies reported here. As such individual PK models were created for animals treated in different protocols. For future pharmacodynamic analyses we will want to include every animal to assess for toxicity. In our study only single daily doses of cefepime were given, thus is unknown whether multiple daily doses demonstrate concentration mediated changes to cerebral cortex and hippocampus transit.

In summary, this data has provided insight on the neurotoxicity threshold. The integrated animal data and PK models may have direct implications for human health outcomes and can provide a framework for optimal treatments regimens, especially in the setting of increasing antimicrobial resistance.

## Acknowledgements

I’d like to thank my mentor, Dr. Marc Scheetz and committee members, Dr. Gwendolyn Pais and Dr. Annette Gilchrist for their expertise and guidance throughout this thesis project as well as lab members Dr. Patti Engel, Dr. Roxane Rohani, Dr. Jack Chang, Dr. Nathaniel Rhodes, Sylwia Marianski, Kimberly Valdez, and Zoe Gibson for their collaboration on the project. I would also like to thank the MWU Animal Facility and Research Core Facility.

## Transparency declarations

None to declare.

